# The CTCF-H3K27me3 axis supports melanoma cell migration by repressing cholesterol biosynthesis

**DOI:** 10.1101/2023.02.23.529650

**Authors:** Lukasz Stanislaw Kaczmarczyk, Nehora Levi, Mali Salmon-Divon, Gabi Gerlitz

**Affiliations:** Department of Molecular Biology, Faculty of Life Sciences and Ariel Center for Applied Cancer Research, Ariel University, Ariel 40700, Israel; Adelson School of Medicine, Ariel University, Ariel 40700, Israel

**Keywords:** SREBP, cholesterol, CTCF, heterochromatin, transcriptional control, cell migration

## Abstract

CTCF is a key factor in three-dimensional chromatin folding and transcriptional control that was found to affect cancer cell migration by a mechanism that is still poorly understood. To identify this mechanism, we used mouse melanoma cells with a partial loss of function (pLoF) of CTCF. We found that CTCF pLoF inhibits cell migration rate while leading to an increase in the expression of multiple enzymes in the cholesterol biosynthesis pathway along with an elevation in the cellular cholesterol levels. Inhibition of cholesterol synthesis in CTCF pLoF cells restored the cellular migration rate, suggesting that CTCF supports cell migration by suppressing cholesterol synthesis. Detailed analysis of the promoter of *Hmgcs1*, an early enzyme in the cholesterol synthesis pathway, revealed that CTCF enables PRC2 recruitment to the promoter and deposition of H3K27me3 to prevent SREBP2 binding and activation of transcription. By this mechanism CTCF fine-tunes cholesterol levels to support cell migration. Notably, genome wide association studies suggest a link between CTCF and cholesterol-associated diseases, thus CTCF emerges as a new regulator of cholesterol biosynthesis.

## Introduction

Cell migration is a fundamental process in embryogenesis and in additional systems such as the immune system [1–3]. Mutations and deregulation of the cellular migration processes are linked to various human diseases including cancer [4]. During cell migration, the nucleus undergoes major changes in its position, structure and morphology [5–10]. One of these changes is global chromatin condensation, which was initially found in mouse melanoma cells migrating in the wound healing assay [11,12] and later on in additional cell types including human breast cancer cells [13], human and mouse CD4^+^ T-cells [14], rat tenocytes [15] and human fibrosarcoma cells [16]. Global chromatin condensation supports cell migration as concluded from dozens of experiments in which interfering with various chromatin condensation factors including histone modifying enzymes, DNA modifying enzymes and histone H1 altered cell migration rate [11,12,14,16,17,17–20]. Heterochromatin supports cell migration by both genetic and mechanical ways [19]. The mechanical properties of the nucleus are affected by chromatin organization: condensed chromatin was shown to enhance nuclear deformability [20] and elasticity [15,21,22] that is thought to enable a bigger forward jump of the nucleus once it is extracted from a narrow pore, leading to an increase in cell migration rate [23]. In addition, in migrating cells the heterochromatin marker H3K27me3 was found to support transcriptional changes in various genes as well as to prevent changes in transcription of hundreds of other genes [24].

To enhance the understanding of the importance of chromatin organization for cancer cell migration we focused on the chromatin architectural protein CCCTC-binding factor (CTCF). CTCF contains unstructured N’ and C’ termini with a central region that contains 11 zinc-finger domains, which are responsible for binding a long and complex DNA motif [25–27]. CTCF is involved in regulation of topologically associating domains (TADs) that are chromatin domains of strong self-contact comprising several hundred kb. TADs are thought to form by a loop-extrusion mechanism in which cohesin complexes form chromatin loops by their ATP dependent molecular motor activity until halted by chromatin bound CTCF; thus, the resulted TAD is folded on itself while it is bound by cohesins and CTCF at its boundaries. Interactions between non-adjacent chromatin elements such as promoter and enhancer more commonly occur within a TAD rather than between separated TADs [27–29]. Loss of CTCF binding at TAD boundaries weakens the boundaries and their ability to insulate contacts between domains leading to transcriptional deregulation [30,31]. CTCF can affect transcription also by direct promoter binding to either repress transcription probably by recruitment of additional repressive factors such as the transcriptional repressor SIN3A [25,32] or to promote transcription by recruitment of transcription factors, repression of antisense transcription [25,32] and promotion of promoter-enhancer interactions [33,34].

CTCF involvement in cell migration is ambiguous; on one hand it was shown to repress breast cancer cell migration and invasion [35,36], while on the other hand it was shown to support migration and/or invasion of various primary and cancerous cells including human corneal epithelial cells [37,38], cortical neurons [39], skin-resident dendritic cells [40], prostate cancer cells [41,42], ovarian cancer cells [43], gastric cancer cells [44] and melanoma cells [45]. Although it is assumed that CTCF affects cell migration by its ability to regulate transcription, the exact mechanism by which CTCF controls transcription to control cell migration has not been determined yet.

Here we show that a partial loss of function (pLoF) of CTCF leads to slower migration of melanoma cells. Transcriptome analysis identified upregulation of cholesterol biosynthesis enzymes along with elevated cholesterol levels in CTCF pLoF cells. The elevated levels of cholesterol in CTCF pLoF cells interfered with their migration rate since inhibition of cholesterol biosynthesis restored their migration capabilities. Detailed analysis of the promoter of the cholesterol biosynthesis enzyme *Hmgcs1* showed that CTCF promotes recruitment of PRC2 to this promoter to methylate H3K27. A methylation process that prevents binding of SREBP2 to the promoter and increased transcription of *Hmgcs1*. Thus, CTCF emerges as a new regulator of cholesterol biosynthesis.

## Results

To evaluate the effect of CTCF on cell migration we used the mouse melanoma B16-F1 cells in which we reduced CTCF expression levels by the CRISPR-Cas9 system [46]. Targeting of Cas9 to the beginning of the *CTCF* gene led to elimination of the full-length form of CTCF and the appearance of truncated forms of CTCF at low levels that are probably formed from downstream AUGs. The total amount of the CTCF shorter forms is 30-50% of the full-length protein in the parental cells. In addition, CTCF truncated forms have lower chromatin binding affinity than the full-length protein, thus these clones have a partial loss of function (pLoF) of CTCF. CTCF pLoF led to 20-40% reduction in the cellular migration rate in the wound healing assay and to 48-86% reduction in the cellular migration rate in the Transwell assay (Fig. 1a-c).

**Fig. 1.**
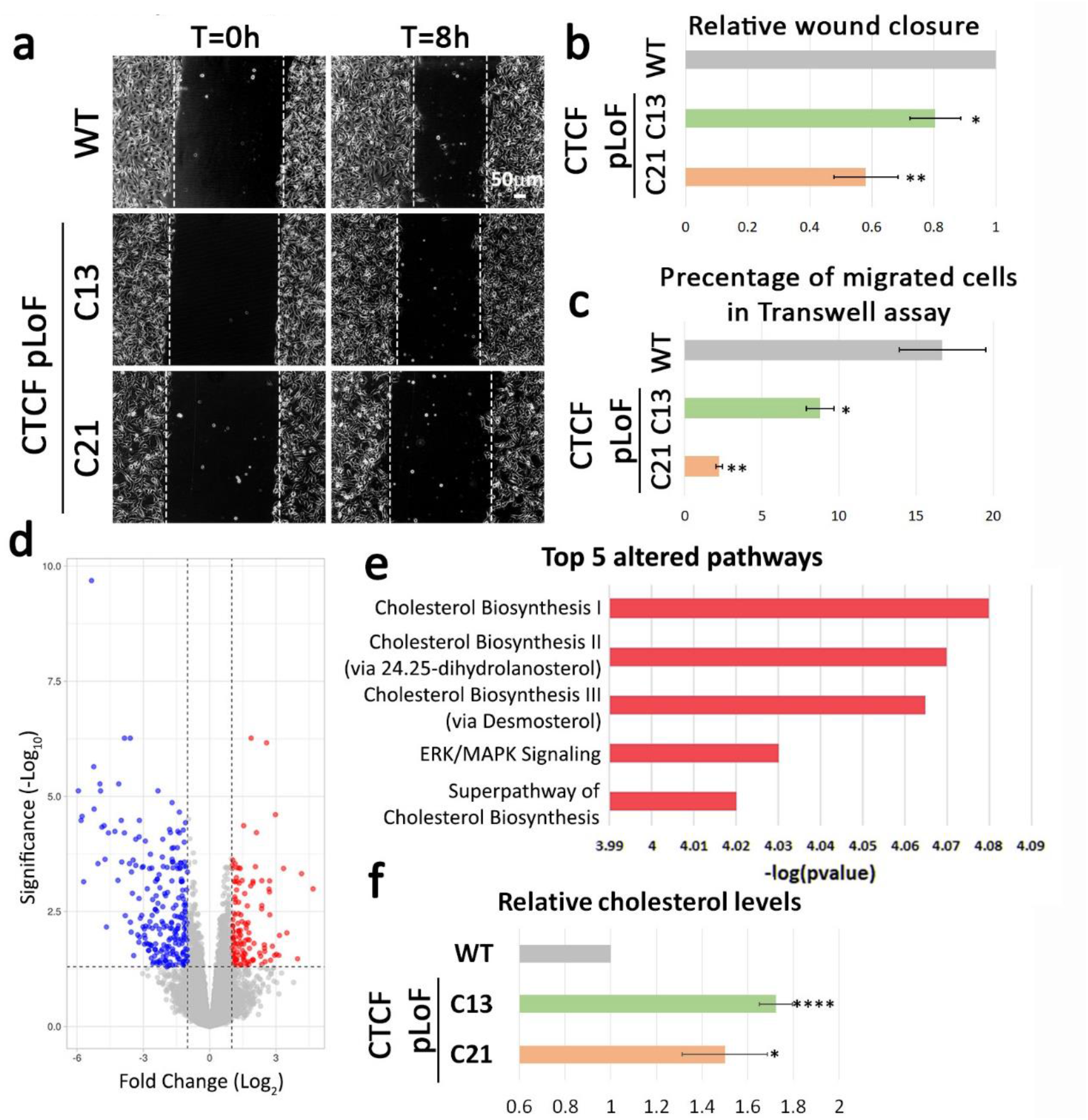
CTCF pLoF attenuates cell migration and elevates cellular cholesterol levels. (**a**) CTCF pLoF attenuates the migration rate of B16-F1 cells in the wound healing assay. Confluent parental B16-F1 cells (WT) and two CTCF pLoF clones (C13 and C21) were induced to migrate for 8h. Representative pictures of the cells immediately after the scratch (0 hours) and eight hours after the scratch are shown. The wound edge is indicated with dashed white line. Scale bar: 50μm. (**b**) For quantification, the area covered by the cells after incubation of eight hours was measured using ImageJ and normalized to the parental B16-F1 cells (WT). The graph shows the mean relative scratch coverage in five independent experiments ± s.e. Statistical significance in comparison to WT cells was calculated by the Student’s *t*-test, \**P*<0.05, \*\**P*<0.01. (**c**) CTCF pLoF attenuates the migration rate of B16-F1 cells in the Transwell assay. Three hours after plating the WT and CTCF pLoF cells on top of 5 μm filters the cells were fixed, permeabilized, and stained with Hoechst reagent. In each experiment the fraction of the cells migrated to the lower side of the filter was calculated. The bar graph represents the mean percentage of migrating cells from the total number of cells on the filter in three independent experiments ± s.e. Statistical significance in comparison to WT cells was calculated by the Student’s *t*-test, \**P*<0.05, \*\**P*<0.01. (**d**) Altered gene expression in CTCF pLoF clones. A volcano plot of altered genes comparing two combined results of two CTCF pLoF clones to WT; four repetitions for each cell type (WT, C13, C21). The graph represents unchanged genes (grey), genes with increased expression (red, 131 genes with fold change ≥2 and *P*<0.05) and genes with decreased expression (blue, 233 genes with fold change ≤-2 and *P* <0.05). (**e**) Top 5 pathways that are altered in CTCF pLoF clones identified by IPA analysis of 364 differentially expressed genes. The top 5 altered pathway were all activated. (**f**) Cellular cholesterol levels are increased in CTCF pLoF cells. The relative cholesterol level WT and CTCF pLoF clones normalized to the total protein level in relation to WT in three independent experiments ± s.e. Statistical significance in comparison to WT cells was calculated by the Student’s *t*-test, \**P*<0.05, \*\*\*\**P*<0.0001.

To look for the molecular mechanism for that phenotype we analysed the transcriptome of CTCF pLoF cells and identified 364 significantly changed genes with at least 2-fold change in their transcript level; 233 genes were downregulated, and 131 genes were upregulated (Fig. 1d, Sup. Fig. 1). To identify affected pathways, we used the Ingenuity Pathway Analysis (IPA, Qiagen). As shown in Fig. 1e, the most significantly up-regulated pathway in CTCF pLoF cells was cholesterol biosynthesis. Cholesterol measurement confirmed that CTCF pLoF clones have an increase of 50-70% in their cellular cholesterol levels (Fig. 1f).

To validate the dependency of reduced cell migration rate on elevated cholesterol levels in CTCF pLoF cells we repeated the migration assays in cells treated with Fatostatin. Fatostatin is an inhibitor of activation of Sterol Regulatory-Element Binding Proteins (SREBPs) that are the master regulators of transcriptional activation of cholesterol biosynthesis enzymes [47,48]. As shown in Fig. 2, Fatostatin treatment elevated the migration rate of CTCF pLoF cells to almost the migration rate of WT cells in both the wound healing assay and the Transwell assay. Thus, it seems that elevated cholesterol levels in CTCF pLoF cells are the main cause for their faulty migration capabilities.

**◄Fig. 2.**
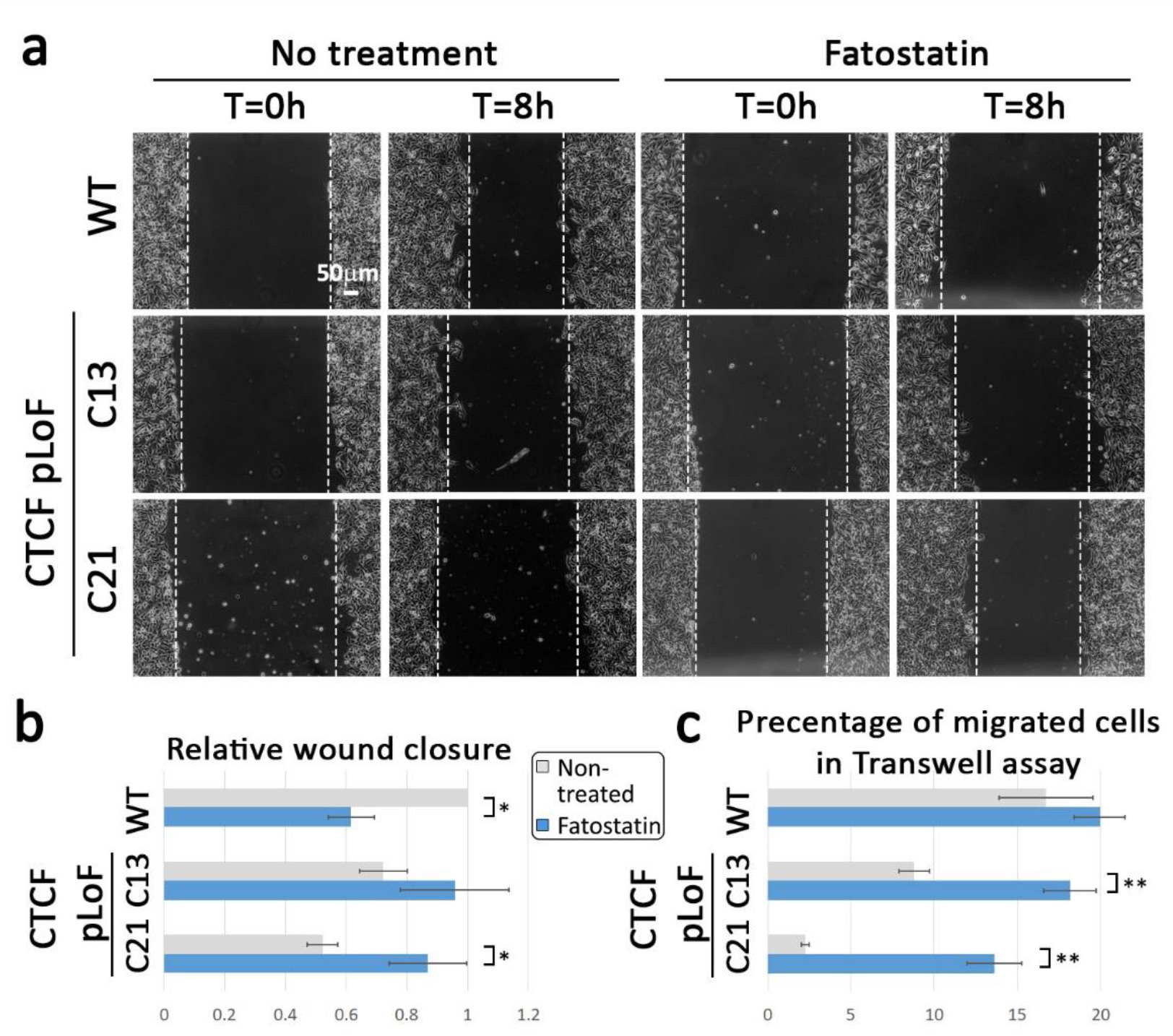
Inhibition of cholesterol synthesis improves the migration capabilities of CTCF pLoF cells. (**a**) Fatostatin improves the migration rate of CTCF pLoF cells in the wound healing assay. Confluent parental B16-F1 cells (WT) and two CTCF pLoF clones (C13 and C21) treated and non-treated with 5 mM Fatostatin for 24 hours prior to the experiment were induced to migrate for 8h. Representative pictures of the cells immediately after the scratch (0 hours) and eight hours after the scratch are shown. The wound edge is indicated with dashed white line. Scale bar: 50mm. (**b**) For quantification, the area covered by the cells after incubation of eight hours was measured using ImageJ and normalized to the non-treated parental B16-F1 cells (WT). The graph shows the mean relative scratch coverage in three independent experiments ± s.e. Statistical significance was calculated by the Student’s *t*-test, \**P*<0.05. (**c**) Fatostatin improves the migration rate of CTCF pLoF cells in the Transwell assay. Parental B16-F1 cells (WT) and two CTCF pLoF clones (C13 and C21) treated and non-treated with 5 mM Fatostatin for 24 hours prior to the experiment were plated on top of 5 μm filters. After three hours the cells were fixed, permeabilized, and stained with Hoechst reagent. In each experiment the fraction of the cells migrated to the lower side of the filter was calculated. The bar graph represents the mean percentage of migrating cells from the total number of cells on the filter in three independent experiments ± s.e. Statistical significance was calculated by the Student’s *t*-test, \*\**P*<0.01.

IPA upstream regulator analysis suggested SREBPs as possible altered factors in CTCF pLoF cells (Sup. Fig. 2). We did not identify any increase in their RNA levels (Fig. 1) nor in their protein levels (Sup. Fig. 2). To analyze SREBPs function we focused on one of the first enzymes in the cholesterol biosynthesis pathway; 3-Hydroxy-3-Methylglutaryl-CoA Synthase 1 (Hmgcs1). First, the dependency of *Hmgcs1* up-regulation in CTCF levels as well as of additional genes was confirmed in a rescue experiment in which mRNA levels of *Hmgcs1* and other altered genes were found to be returned to normal upon over-expression of GFP-fused CTCF (Sup. Fig. 3). Next, we looked for binding of CTCF to the promoter of *Hmgcs1* using CTCF ChIP-seq data from B16-F1 cells [46], but did not find a direct binding of CTCF to the promoter (Sup. Fig. 4). To understand the changes in the function of this promoter we looked for the binding of SREBP2 to this promoter by a ChIP analysis and revealed an increased binding of SREBP2 to *Hmgcs1* promoter in CTCF pLoF cells in comparison to the parental cells (Fig. 3). We hypothesized that Srebp2 binding to *Hmgcs1* promoter is inhibited in the parental cells due to methylation at H3K27, since in a previous study we found that inhibition of H3K27 methylation in B16-F1 cells led to increased expression of cholesterol biosynthesis enzymes [24]. ChIP analysis of *Hmgcs1* promoter revealed reduced H3K27me3 levels in CTCF pLoF cells in comparison to the WT cells. To verify this, we also checked the accumulation of PRC2 that methylates H3K27 at this promoter and indeed binding of Suz12 (one of the subunits in PRC2) was reduced in CTCF pLoF cells (Fig. 3).

**Fig. 3.**
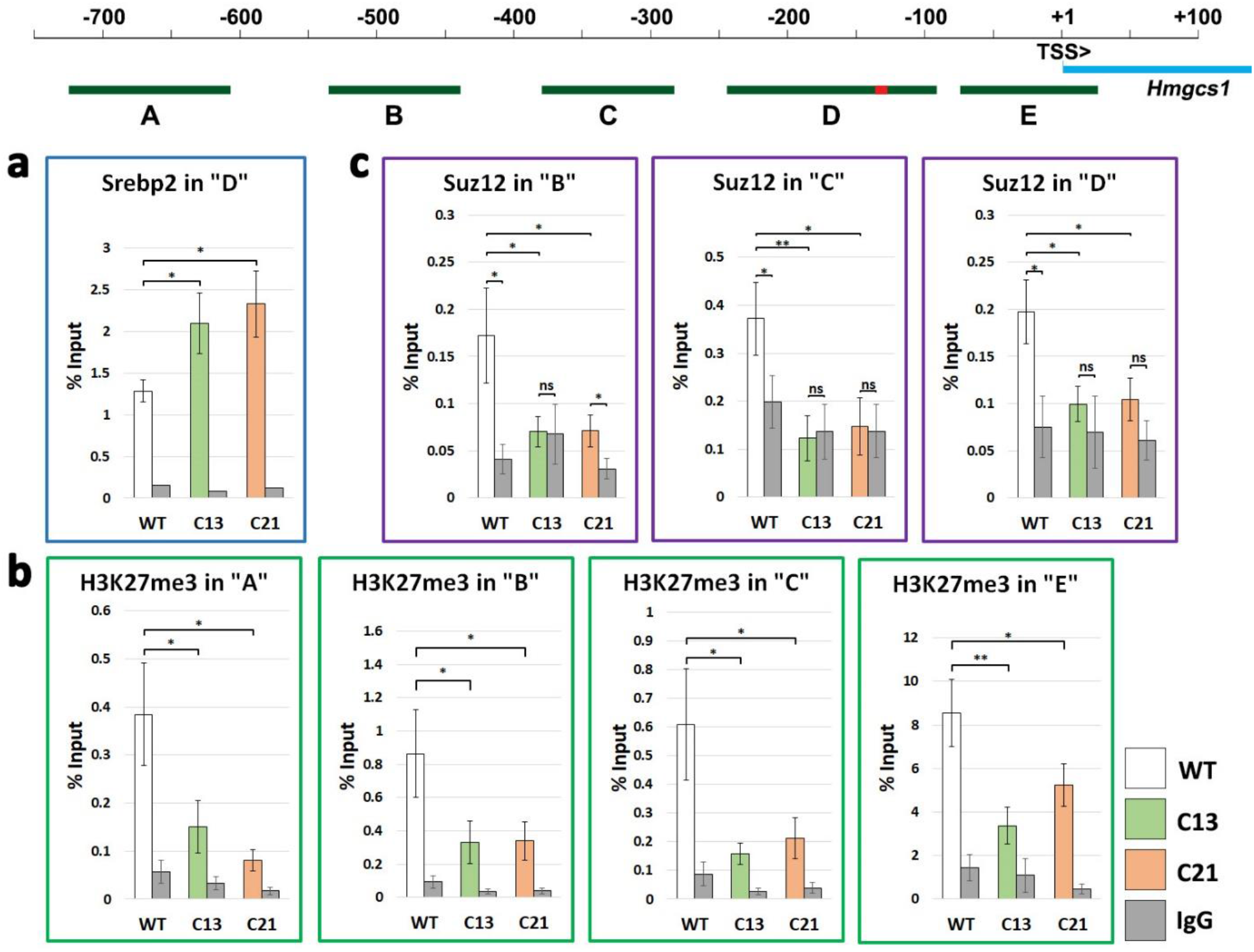
Altered chromatin state of *Hmgcs1* promoter in CTCF pLoF cells. (**a**) A diagram of *Hmgcs1* promoter. SREBP motif is marked by red rectangle. Transcription start site is marked as TSS. Regions A-E were covered by five sets of primers used for ChIP-qPCR. (**b-d**) Mean Srebp2, H3K27me3 and Suz12 levels in the indicated parts of *Hmgcs1* promoter determined by ChIP-qPCR and presented as percentage of input in WT cells (white bars) and in CTCF pLoF clones C13 (green bars) and C21 (orange bars) ± s.e. of six independent experiments. Normal rabbit IgG were used as a negative control to estimate non-specific binding of primary antibodies (grey bars). Statistical significance was calculated by the Student’s *t*-test, \**P*<0.05, \*\**P*<0.01.

**Fig. 4.**
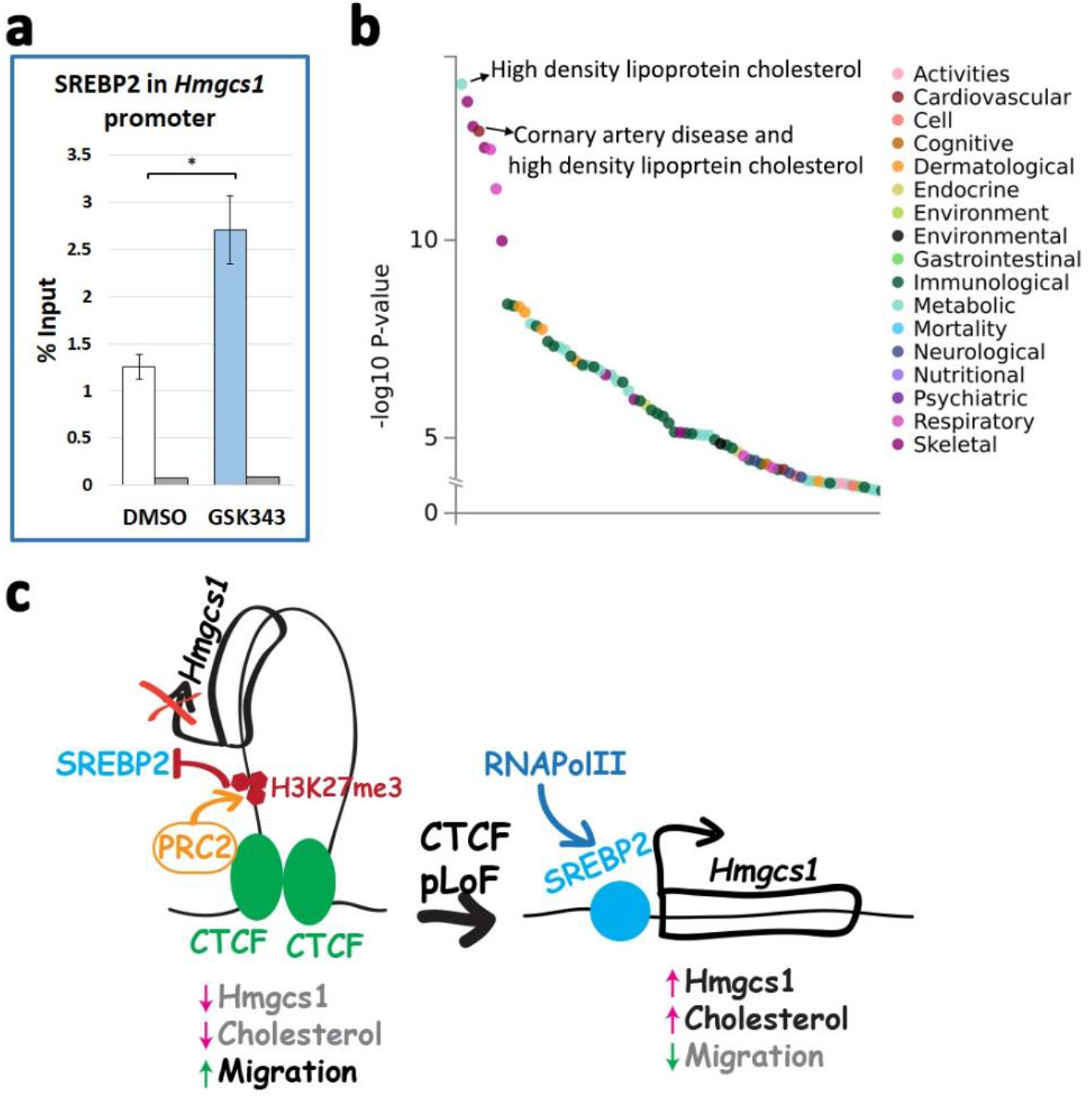
H3K27 methylation prevents Srebp2 binding to *Hmgcs1* promoter and CTCF association with cholesterol-associated diseases. (**a**) Srebp2 binding to *Hmgcs1* promoter was determined in B16-F1 WT cells after 24 hours of GSK343 treatment (blue bar) and in control cells treated with DMSO (white bar) and expressed as a mean percentage of input ± s.e. of six independent experiments. Normal rabbit IgG were used as a control to estimate non-specific binding of target primary antibodies (grey bars). Statistical significance was calculated by the Student’s *t*-test, \**P*<0.05. (**b**) Phenome-wide association between CTCF variants and phenotypes across 4756 GWASs by GWAS-ATLAS (https://atlas.ctglab.nl/). (**c**) A model for the transcriptional control of the cholesterol biosynthesis enzyme Hmgcs1 by CTCF that recruits the H3K27 methyltransferase PRC2 to elevate H3K27me3 at the promoter to prevent the binding of Srebp2.

To prove that H3K27 methylation prevents SREBP2 binding to this promoter we repeated the ChIP analysis for SREBP2 in cells treated with a specific inhibitor for Ezh2, which is the catalytic subunit in PRC2 (GSK343). As shown in Fig. 4, GSK343 treatment led to increased SREBP2 binding to *Hmgcs1* promoter. Interestingly, genome wide association studies (GWAS) query for the association of CTCF with human diseases revealed that the highest correlated disease with CTCF alterations was high density lipoprotein cholesterol (Fig. 4b). Thus, suggesting CTCF as an unknown regulator of cholesterol homeostasis.

## Discussion

Our results suggest that CTCF is crucial for correct cell migration due to its repressive effect on the cholesterol biosynthesis pathway. Reduced levels of CTCF led to impaired migration of the cells in parallel to increased RNA levels of cholesterol biosynthesis enzymes and elevated cellular cholesterol levels. Inhibition of cholesterol synthesis in CTCF pLoF cells alleviated their impaired migration rate. Cholesterol was shown to affect cell migration capabilities by changing membrane mechanical properties such as rigidity as well as by altering the function of membranal proteins and the activation of signal transduction pathways [49]. Detailed analysis of the promoter of *Hmgcs1* that encodes a cholesterol biosynthesis enzyme revealed that CTCF is not bound directly to it, but it is important for recruitment of PRC2 to this promoter to methylate H3K27. This methylation prevents the binding of SREBP2 to this promoter, and thus suppresses transcription of *Hmgcs1* and cholesterol biosynthesis (Fig. 4c). The exact mechanism by which CTCF recruits PRC2 to this promoter is still an open question; CTCF was shown to be able to bind PRC2 [50], thus CTCF may recruit PRC2 to a distant chromatin position from which it spreads towards the promoter. Another option is CTCF-dependent specific 3D contacts of the promoter that either generate a contact with a chromatin site to which PRC2 is already bound or/and prevent the generation of a contact with an active enhancer. Either way, CTCF appears as a metabolic regulator of cholesterol biosynthesis by its ability to modulate PRC2 activity.

Our findings may well be relevant to both high cholesterol-linked diseases as well as cancer treatment. The importance of CTCF for the transcriptional regulation of cholesterol biosynthesis enzymes and the significant genome wide association between CTCF polymorphism, high density lipoprotein cholesterol diseases and coronary artery disease suggest a pathophysiological connection between CTCF and cholesterol-linked diseases. Thus, CTCF polymorphism may be an important factor in the genetic background that promotes high cholesterol. In recent years there are many attempts to treat cancer patients with cholesterol biosynthesis inhibitors such as statins [51,52]. Considering the central role of cell migration in metastasis formation and the inhibitory effect of increased cholesterol levels on cell migration, trying to reduce cholesterol levels in cancer patients that suffer from tumor cells with higher cholesterol levels than the normal levels may not inhibit but rather accelerate the migration rate of these tumor cells and their ability to metastasize.

## Methods

### Cell culture

Mouse melanoma cells B16-F1 were purchased from the ATCC (CRL-6323, ATCC). Parental and CTCF pLoF clones C13 and C21 [46] were grown in DMEM (01-056-1A, Biological industries) supplemented with 10% Fetal Bovine Serum (FBS, 04-007-1A, Biological Industries, Israel), 0.5% penicillin-streptomycin antibiotic mix (03-031-1B, Biological Industries), and 1% L-glutamine (03-020-1B, Biological Industries, Israel) at 37°C, 7% CO_2_. Inhibitors used were and 3 μM of GSK343 (SML0766, Sigma-Aldrich, Rehovot, Israel) for 24 hours. For the rescue experiment CTCF pLoF clones C13 and C21 were transfected with either pKS004-pCAGGS-3XFlag-CTCF-eGFP plasmid (a gift from Elphege Nora, Addgene plasmid #156438) [53] or GFP-NLS (a plasmid expressing GFP fused to a nuclear localization signal that was a gift from Michael Bustin) [21]. Transfection was done with NanoJuice Core Transfection Reagent (71900, Merck) following manufacturer’s instructions with an incubation of 48 h.

### Motility assays

For the wound-healing assay, cells plated in fibronectin-coated 12-well plates were grown to confluence. Then the cells were scratched with the edge of a 10 μl tip, washed three times with DMEM and incubated in DMEM supplemented with 0.5% FCS. When indicated 5μM of Fatostatin (13562, Cayman chemical, Ann Arbor, Michigan, USA) was added 24 hours prior to the migration assay and was kept during the assay. Images of the same fields were collected with an Olympus SC100 camera mounted on an Olympus CKX41 microscope at two time points: immediately after the scratch and eight hours after the scratch. The area covered by the migrating cells was calculated using ImageJ/Fiji by comparison of the same fields between 0 hours and 8 hours. In each experiment four different fields were analyzed for each treatment (two in one well and another two in a second well).

The Transwell® assay was carried out in plate filters with pores diameter of 5 μm (3421, Corning Incorporated, Corning, NY, USA) as previously described [11]. Where indicated Fatostatin was added to the medium as in the wound healing assay. For counting purposes, the cells nuclei were stained with Hoechst reagent.

### Transcriptome analysis

Total RNA was purified by the NucleoZOL kit (740404.200, MACHEREY-NAGEL, Duren, Germany) according to the manufacturer’s instructions. For NGS sequencing by Cel-Seq2 [54] purified RNA from four repetitions for each cell type (WT, C13 and C21) was sent for poly A containing mRNA selection, library preparation and sequencing at the Technion Genome Center. Fastq files were deposited in SRA (BioProject ID: PRJNA935672). For RT-qPCR 0.4μg of RNA was used to prepare cDNA with qPCRBIO cDNA Synthesis Kit (PB30.11-10, PCR Biosystems, London, UK). qPCR was done with Fast SYBR Green Master Mix (4385612, Thermo Fisher Scientific, Vilnius, Lithuania). The various genes were normalized to the RNA levels of the ribosomal protein Rps29. Primers used: *Rps29*: forward: 5’-tcgttgggcgtctgaaggcaa-3’, reverse: 5’-cggaagcactggcggcacat-3’; *Eva1a*: forward: 5’-ttgggaatggctctgctc-3’, reverse: 5’-ggagatcctcatcaccaagg-3’; *Hs3st1*: forward: 5’-gtgtgaatttgctccaaagg-3’, reverse: 5’-ag-caaggctccctcttgatt-3’; *Pou3f1*: forward: 5’-aacaagtggctggaggagac-3’, reverse: 5’-ggcttgggacactt-gagaaa-3’; *Nov*: forward: 5’-tcgtctctgcatcgttcgg-3’, reverse: 5’-catcactgcagaccccaca-3’; *Pgbd5*: forward: 5’-tacaaggtccagcccttcct-3’, reverse: 5’-tttccgctttttcctcttcc-3’; *Hsd17b7*: forward: 5’-ggacgttgctcctacccata-3’, reverse: 5’-cgcgtatttggtcagaggat-3’; *Hmgcs1*: Forward: 5’-cctggac-cgctgctattc-3’, reverse: 5’-ttcaggaacatccgagctaga; *Lgals3*: forward: 5’-agacagtcagccttcccct-3’, re-verse: 5’-agttggctgatttcccggag-3’; *Egln3:* forward: 5’-caggttatgttcgccatgtg-3’, reverse: 5’-ggctccacgtctgctacaa-3’; *Rgs16:* forward: 5’-cctgcgaggagttcaagaag-3’, reverse: 5’-tggtagtgg-cagcttgtagg-3’; *Selenbp1:* forward: 5’-ctaacccacagaagccccg-3’, reverse: 5’-gctggatcatctgagggcc-3’; *Col1a2*: forward: 5’-cccctggtcttactgggaac-3’, reverse: 5’-caggtccttggaaaccttga-3’; *Mef2c*: forward: 5’-ggggactatggggagaaaaa-3’, reverse: 5’-atctcacagtcgcacagcac-3’; *Dek:* forward: 5’-cag-tgacacaagggaagggt-3’, reverse: 5’-ccactgaactgacccacgt-3’; *Ddit4:* forward: 5’-accag-ttcgctcacccttc-3’, reverse: 5’-aaacgatcccagaggctagg-3’; *Slc2a1:* forward: 5’-ttgtt-gtagagcgagctgga-3’, reverse: 5’-aaggccacaaagccaaagat-3’; *Egln1:* forward: 5’-cctggatcgaggg-caaaga-3’, reverse: 5’-tagcctgttccgttgcctg-3’; *Frs3*: forward: 5’-gtgttcgagggcagaggac-3’, reverse: 5’-cagagaccgtgtagccagc-3’; *Ctgf:* forward: 5’-cctgcgacccacacaagg-3’, reverse: 5’-gctgcttt-ggaaggactcac-3’; *Celsr3:* forward: 5’-caacaagccccggacagat-3’, reverse: 5’-cagctgcacacgagtagttg-3’; *Mgarp:* forward: 5’-tggaacatctggctccaata-3’, reverse: 5’-caactccgcctttgtttgtt-3’; *Runx2:* 5’-tgagatttgtgggccggag-3’, reverse: 5’-agcttctgtctgtgccttct-3’; *Ttbk1*: 5’-gcttcaaggcaaggaccac-3’, 5’-ccaggatctgcttgcccag-3’.

### Cholesterol assay

The cholesterol assay was carried out according to the manufacturer protocol using the Cholesterol/Cholesterol Ester Assay Kit (ab65359, Abcam, Cambridge, UK) and normalized to total protein content as determined by Bradford Assay.

### ChIP

Chromatin immunoprecipitation (ChIP) was performed as described previously [24] with some modifications: for each sample the lysate from 10^7^ cells (total ∼700μl) was divided into three fractions: sample (330μl), IgG control (330μl) and input (30μl). Samples were added to Magna ChIP™ Protein A+G Magnetic Beads (16-663, Sigma-Aldrich) precoated with specific antibody: 4μg of rabbit anti-Srebp2 (ab112046, Abcam, Cambridge, UK) or 4μg rabbit anti-H3K27me3 (07-449, Sigma-Aldrich) or 3μl of rabbit anti-Suz12 [3737, Cell Signaling, Danvers, MA, USA). IgG controls were added to magnetic beads precoated with 4μg of normal rabbit IgG (sc2027, Santa Cruz Biotechnology, Dallas, TX, USA). Quantitative PCR was carried with specific primers for the regions A-E in the promoter of *Hmgcs1*: (A) forward: 5’-tggaccaag tgccaaaataa-3’, reverse: 5’-gaatgatgcagggcttgtct-3’; (B) forward: 5’-gccaaggtagtttaaattggaca-3’, reverse: 5’-gcaaattgcatctgcatcac-3’; (C) forward: 5’-tacaggtgaagccgagctgt-3’ or 5’-gaagccgagctgtggttg-3’, reverse: 5’-actcctggggagagatggtc-3’; (D) forward: 5’-gaccttcaattggtcggaga-3’, reverse: 5’-ggtaaggggtgggaacaaag-3’; (E) forward: 5’-agggaaaaccctagcgagtc-3’, reverse: 5’-cggctac-ctctgccaatc-3’.

### Western blot analysis

Whole cell lysate was prepared by incubation of cells with 2x Sample Buffer at 96°C for 10min and stored at -20°C. Antibodies used: mouse anti-Srebp1 (39939, Active Motif, Carlsbad, CA, USA, dilution 1:1,000), rabbit anti-Srebp2 (112046, Abcam, Cambridge, UK, dilution 1:1,000), mouse anti-β-actin (A1978, Sigma-Aldrich, dilution 1:5,000).

### Bioinformatic analysis

Sequencing reads from four replicates of RNA-seq samples for each pLoF clone and WT were aligned to the mouse genome (mm10) by Hisat2 program [55,56] after removal of adapter sequences and critical examination of quality controls using Trim Galore and FastQC respectively with default parameters. Next, the number of reads mapping each mouse gene (as annotated in Ensembl release GRCm38.100) was counted using the featureCounts program [57]. Differential expression analysis was performed using edgeR and Limma packages [58,59] from the Bioconductor framework [60]. Features with less than 1 read per million in four samples were removed. The remaining gene counts were normalized using the TMM method, followed by voom transformation [61]. Linear models as implemented by the Limma package were used to find differentially expressed genes between the two CTCF pLoF clones and the WT cells. Obtained lists with significantly changed genes (*p*<0.05, fold-change ≥2) were used in Ingenuity Pathway Analysis (IPA, Qiagen).

## Author contributions

Conceptualization, L.S.K., N.L., M.S.D. and G.G.; Methodology, L.S.K. and N.L.; Investigation, L.S.K. and N.L.; Formal Analysis -L.S.K., N.L. and M.S.D.; Writing -Original Draft, G.G.; Writing -Review & Editing, L.S.K., N.L., M.S.D. and G.G., Funding Acquisition, M.S.D. and G.G.

## Supplementary Figures

**Sup. Fig. 1.**
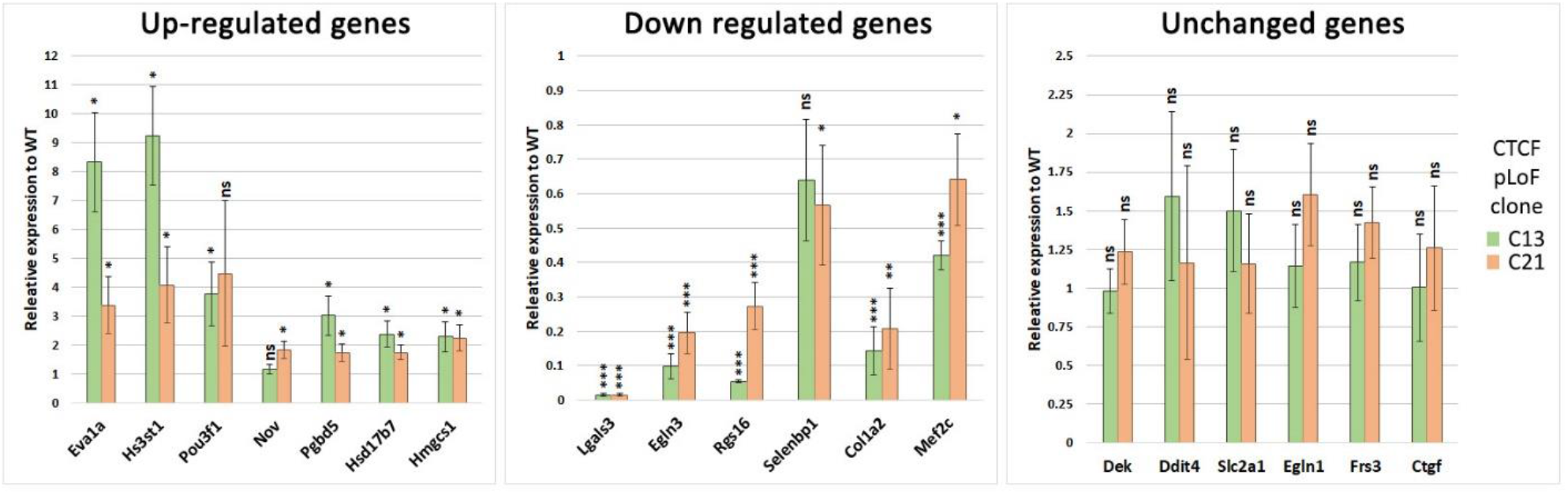
Verification of RNA-seq based transcriptome profiling in CTCF pLoF cells by RT-qPCR. Relative RNA levels of up-regulated, down regulated and unchanged genes in the RNA-seq analysis were determined by RT-PCR. Graph bars represent means of ratios between RNA levels in CTCF pLoF cells and WT cells ± s.e. of three independent experiments. Statistical significance was calculated by the Student’s *t*-test, \**P*<0.05.

**Sup. Fig. 2.**
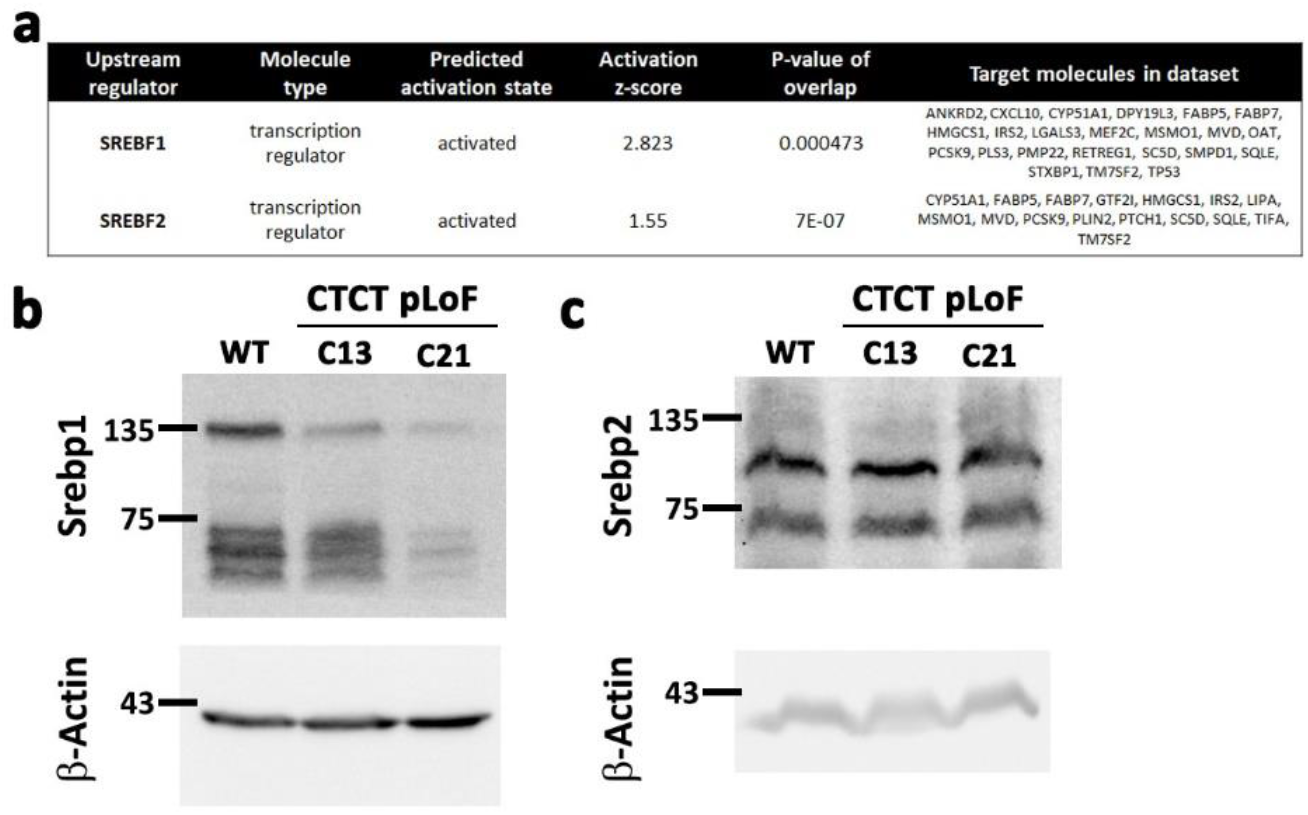
SREBPs in CTCF pLoF cells. (**a**) Predicted upstream regulators by IPA analysis based on 364 differentially expressed genes in CTCF pLoF cells identified SREBPs (SREBFs) as possible activated transcription factors in these cells. (**b-c**) Protein levels of Srebp1 and Srebp2 in CTCF pLoF cells as determined by Western blot analyses.

**Sup. Fig. 3.**
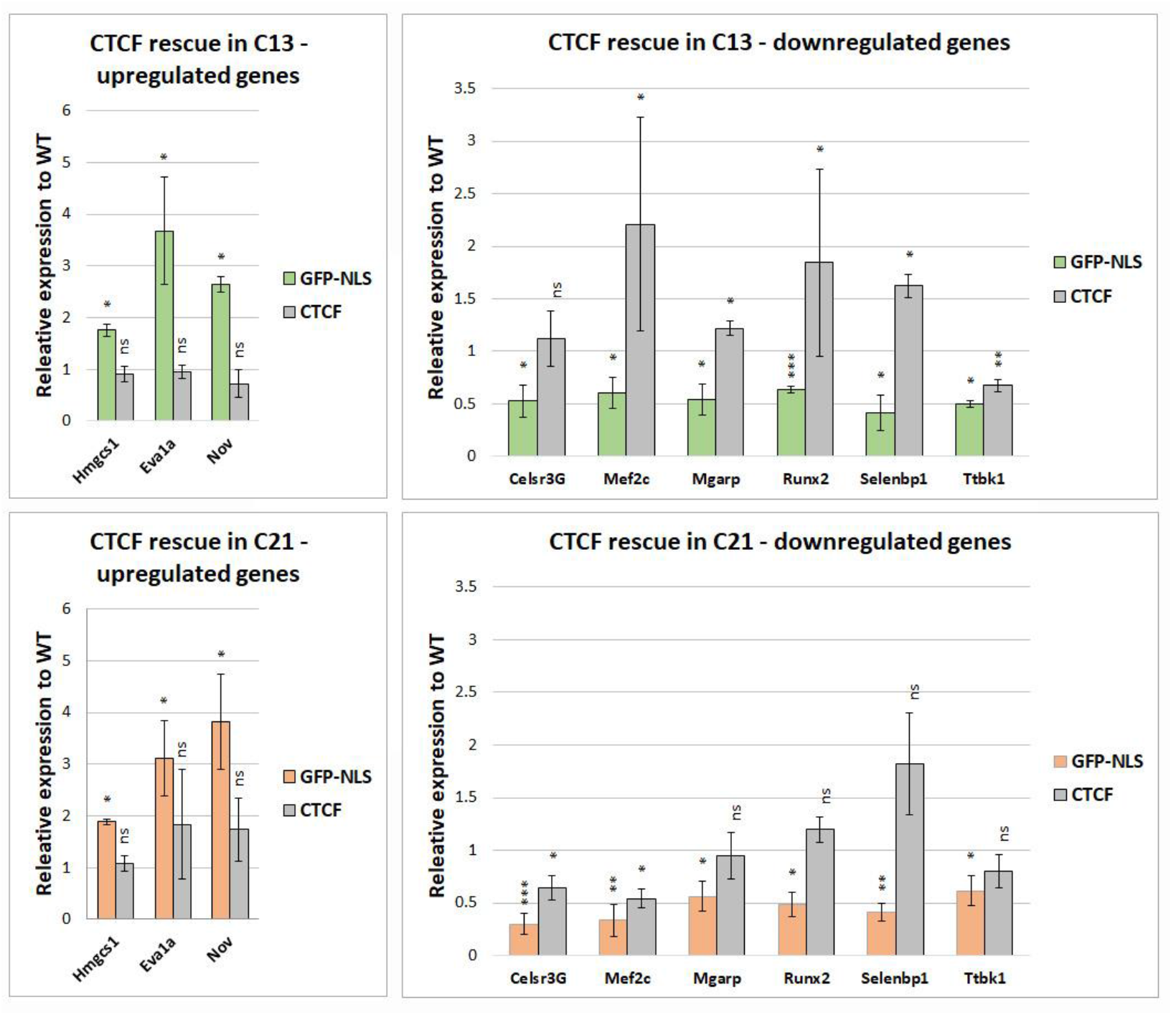
Overexpression of CTCF reverts the expression levels of various genes in CTCF pLoF cells. Relative RNA levels of downregulated and upregulated genes in CTCF pLoF clones C13 (upper panels) and C21 (lower panels) after overexpression of CTCF or GFP-NLS that was served as a negative control. Results are presented as relative expression level in comparison to WT cells ± s.e. of three independent experiments. Statistical significance was calculated by the Student’s *t*-test, \**P*<0.05, \*\**P*<0.01, \*\*\**P*<0.001.

**Sup. Fig. 4.**
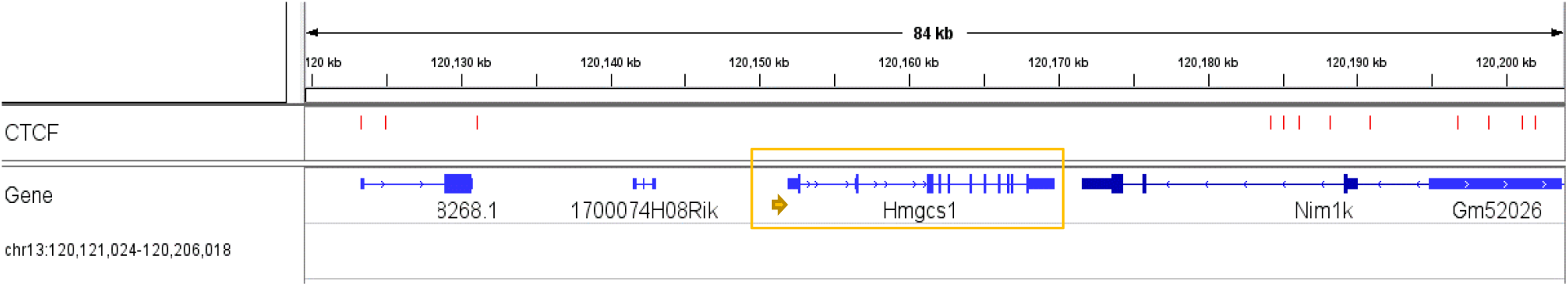
CTCF binding around *Hmgcs1*. Integrative Genomic Viewer (IGV) browser view of CTCF peaks identified by ChIP-seq (red dashes) around *Hmgcs1* (marked with an orange rectangle). The promoter of *Hmgcs1* is indicated with an orange arrow.

## References

1. Blackley DG, Cooper JH, Pokorska P, Ratheesh A. 2021 Mechanics of developmental migration. Seminars in Cell & Developmental Biology 120, 66–74. (doi:10.1016/j.semcdb.2021.07.002)

2. Sapir T, Sela-Donenfeld D, Karlinski M, Reiner O. 2022 Brain Organization and Human Diseases. Cells 11, 1642. (doi:10.3390/cells11101642)

3. Delgado M, Lennon-Duménil A-M. 2022 How cell migration helps immune sentinels. Front. Cell Dev. Biol. 10, 932472. (doi:10.3389/fcell.2022.932472)

4. Novikov NM, Zolotaryova SY, Gautreau AM, Denisov EV. 2021 Mutational drivers of cancer cell migration and invasion. Br J Cancer 124, 102–114. (doi:10.1038/s41416-020-01149-0)

5. Wolf K, Wu YI, Liu Y, Geiger J, Tam E, Overall C, Stack MS, Friedl P. 2007 Multi-step pericellular proteolysis controls the transition from individual to collective cancer cell invasion. Nat Cell Biol 9, 893–904. (doi:10.1038/ncb1616)

6. Wolf K et al. 2013 Physical limits of cell migration: Control by ECM space and nuclear deformation and tuning by proteolysis and traction force. The Journal of Cell Biology 201, 1069–1084. (doi:10.1083/jcb.201210152)

7. Friedl P, Wolf K. 2009 Proteolytic interstitial cell migration: a five-step process. Cancer Metastasis Rev. 28, 129–135. (doi:10.1007/s10555-008-9174-3)

8. McGregor AL, Hsia C-R, Lammerding J. 2016 Squish and squeeze—the nucleus as a physical barrier during migration in confined environments. Current Opinion in Cell Biology 40, 32–40. (doi:10.1016/j.ceb.2016.01.011)

9. Lele TP, Dickinson RB, Gundersen GG. 2018 Mechanical principles of nuclear shaping and positioning. J. Cell Biol. 217, 3330–3342. (doi:10.1083/jcb.201804052)

10. Yamada KM, Sixt M. 2019 Mechanisms of 3D cell migration. Nat Rev Mol Cell Biol 20, 738–752. (doi:10.1038/s41580-019-0172-9)

11. Gerlitz G, Livnat I, Ziv C, Yarden O, Bustin M, Reiner O. 2007 Migration cues induce chromatin alterations. Traffic 8, 1521–1529. (doi:10.1111/j.1600-0854.2007.00638.x)

12. Gerlitz G, Bustin M. 2010 Efficient cell migration requires global chromatin condensation. J. Cell. Sci. 123, 2207–2217. (doi:10.1242/jcs.058271)

13. Tanaka H, Nishioka Y, Yokoyama Y, Higashiyama S, Matsuura N, Matsuura S, Hieda M. 2012 Nuclear envelope-localized EGF family protein amphiregulin activates breast cancer cell migration in an EGF-like domain independent manner. Biochem Biophys Res Commun 420, 721–6. (doi:10.1016/j.bbrc.2012.03.045)

14. Zhang X, Cook PC, Zindy E, Williams CJ, Jowitt TA, Streuli CH, MacDonald AS, Redondo-Muñoz J. 2016 Integrin α4β1 controls G9a activity that regulates epigenetic changes and nuclear properties required for lymphocyte migration. Nucleic Acids Research 44, 3031–3044. (doi:10.1093/nar/gkv1348)

15. Zhang B, Luo Q, Chen Z, Shi Y, Ju Y, Yang L, Song G. 2016 Increased nuclear stiffness via FAK-ERK1/2 signaling is necessary for synthetic mechano-growth factor E peptide-induced tenocyte migration. Scientific Reports 6, 18809. (doi:10.1038/srep18809)

16. Hsia C-R, McAllister J, Hasan O, Judd J, Lee S, Agrawal R, Chang C-Y, Soloway P, Lammerding J. 2022 Confined migration induces heterochromatin formation and alters chromatin accessibility. iScience 25, 104978. (doi:10.1016/j.isci.2022.104978)

17. Maizels Y, Elbaz A, Hernandez-Vicens R, Sandrusy O, Rosenberg A, Gerlitz G. 2017 Increased chromatin plasticity supports enhanced metastatic potential of mouse melanoma cells. Exp. Cell Res. 357, 282–290. (doi:10.1016/j.yexcr.2017.05.025)

18. Lu Z et al. 2020 Epigenetic therapy inhibits metastases by disrupting premetastatic niches. Nature 579, 284–290. (doi:10.1038/s41586-020-2054-x)

19. Gerlitz G. 2020 The Emerging Roles of Heterochromatin in Cell Migration. Front. Cell Dev. Biol. 8, 394. (doi:10.3389/fcell.2020.00394)

20. Madrazo E, González-Novo R, Ortiz-Placín C, García de Lacoba M, González-Murillo Á, Ramírez M, Redondo-Muñoz J. 2022 Fast H3K9 methylation promoted by CXCL12 contributes to nuclear changes and invasiveness of T-acute lymphoblastic leukemia cells. Oncogene 41, 1324–1336. (doi:10.1038/s41388-021-02168-8)

21. Furusawa T, Rochman M, Taher L, Dimitriadis EK, Nagashima K, Anderson S, Bustin M. 2015 Chromatin decompaction by the nucleosomal binding protein HMGN5 impairs nuclear sturdiness. Nat Commun 6, 6138. (doi:10.1038/ncomms7138)

22. Strom AR et al. 2021 HP1α is a chromatin crosslinker that controls nuclear and mitotic chromosome mechanics. eLife 10, e63972. (doi:10.7554/eLife.63972)

23. Krause M et al. 2019 Cell migration through three-dimensional confining pores: speed accelerations by deformation and recoil of the nucleus. Phil. Trans. R. Soc. B 374, 20180225. (doi:10.1098/rstb.2018.0225)

24. Segal T, Salmon-Divon M, Gerlitz G. 2018 The Heterochromatin Landscape in Migrating Cells and the Importance of H3K27me3 for Associated Transcriptome Alterations. Cells 7, 205. (doi:10.3390/cells7110205)

25. Braccioli L, de Wit E. 2019 CTCF: a Swiss-army knife for genome organization and transcription regulation. Essays in Biochemistry 63, 157–165. (doi:10.1042/EBC20180069)

26. Wu Q, Liu P, Wang L. 2020 Many facades of CTCF unified by its coding for three-dimensional genome architecture. Journal of Genetics and Genomics 47, 407–424. (doi:10.1016/j.jgg.2020.06.008)

27. Dehingia B, Milewska M, Janowski M, Pękowska A. 2022 CTCF shapes chromatin structure and gene expression in health and disease. EMBO Reports 23. (doi:10.15252/embr.202255146)

28. Lazniewski M, Dawson WK, Rusek AM, Plewczynski D. 2019 One protein to rule them all: The role of CCCTC-binding factor in shaping human genome in health and disease. Semin Cell Dev Biol 90, 114–127. (doi:10.1016/j.semcdb.2018.08.003)

29. Misteli T. 2020 The Self-Organizing Genome: Principles of Genome Architecture and Function. Cell 183, 28–45. (doi:10.1016/j.cell.2020.09.014)

30. Nora EP, Goloborodko A, Valton A-L, Gibcus JH, Uebersohn A, Abdennur N, Dekker J, Mirny LA, Bruneau BG. 2017 Targeted Degradation of CTCF Decouples Local Insulation of Chromosome Domains from Genomic Compartmentalization. Cell 169, 930–944.e22. (doi:10.1016/j.cell.2017.05.004)

31. Wutz G et al. 2017 Topologically associating domains and chromatin loops depend on cohesin and are regulated by CTCF, WAPL, and PDS5 proteins. EMBO J 36, 3573–3599. (doi:10.15252/embj.201798004)

32. Sun X, Zhang J, Cao C. 2022 CTCF and Its Partners: Shaper of 3D Genome during Development. Genes 13, 1383. (doi:10.3390/genes13081383)

33. Ren G, Jin W, Cui K, Rodrigez J, Hu G, Zhang Z, Larson DR, Zhao K. 2017 CTCF-Mediated Enhancer-Promoter Interaction Is a Critical Regulator of Cell-to-Cell Variation of Gene Expression. Molecular Cell 67, 1049–1058.e6. (doi:10.1016/j.molcel.2017.08.026)

34. Kubo N et al. 2021 Promoter-proximal CTCF binding promotes distal enhancer-dependent gene activation. Nat Struct Mol Biol 28, 152–161. (doi:10.1038/s41594-020-00539-5)

35. Wu J et al. 2017 CCCTC-binding factor inhibits breast cancer cell proliferation and metastasis via inactivation of the nuclear factor-kappaB pathway. Oncotarget 8, 93516–93529. (doi:10.18632/oncotarget.18977)

36. Lebeau B et al. 2022 3D chromatin remodeling potentiates transcriptional programs driving cell invasion. Proc Natl Acad Sci U S A 119, e2203452119. (doi:10.1073/pnas.2203452119)

37. Wang L, Deng SX, Lu L. 2012 Role of CTCF in EGF-Induced Migration of Immortalized Human Corneal Epithelial Cells. Invest. Ophthalmol. Vis. Sci. 53, 946. (doi:10.1167/iovs.11-8747)

38. Wang L, Wu X, Shi T, Lu L. 2013 Epidermal Growth Factor (EGF)-induced Corneal Epithelial Wound Healing through Nuclear Factor κB Subtype-regulated CCCTC Binding Factor (CTCF) Activation. Journal of Biological Chemistry 288, 24363–24371. (doi:10.1074/jbc.M113.458141)

39. Elbert A, Vogt D, Watson A, Levy M, Jiang Y, Brûlé E, Rowland ME, Rubenstein J, Bérubé NG. 2019 CTCF Governs the Identity and Migration of MGE-Derived Cortical Interneurons. J. Neurosci. 39, 177–192. (doi:10.1523/JNEUROSCI.3496-17.2018)

40. Kim T-G et al. 2015 CCCTC-binding factor controls the homeostatic maintenance and migration of Langerhans cells. J Allergy Clin Immunol 136, 713–724. (doi:10.1016/j.jaci.2015.03.033)

41. Shan Z et al. 2019 CTCF regulates the FoxO signaling pathway to affect the progression of prostate cancer. J Cell Mol Med 23, 3130–3139. (doi:10.1111/jcmm.14138)

42. Xing Z, Li S, Liu Z, Zhang C, Bai Z. 2020 CTCF-induced upregulation of HOXA11-AS facilitates cell proliferation and migration by targeting miR-518b/ACTN4 axis in prostate cancer. Prostate 80, 388–398. (doi:10.1002/pros.23953)

43. Zhao L, Yang Y, Yin S, Yang T, Luo J, Xie R, Long H, Jiang L, Zhu B. 2017 CTCF promotes epithelial ovarian cancer metastasis by broadly controlling the expression of metastasis-associated genes. Oncotarget 8, 62217–62230. (doi:10.18632/oncotarget.19216)

44. Liu C et al. 2021 Zinc Finger Protein CTCF Regulates Extracellular Matrix (ECM)-Related Gene Expression Associated With the Wnt Signaling Pathway in Gastric Cancer. Front. Oncol. 10, 625633. (doi:10.3389/fonc.2020.625633)

45. Wen X et al. 2020 An Artificial CTCF Peptide Triggers Efficient Therapeutic Efficacy in Ocular Melanoma. Mol Ther Oncolytics 18, 317–325. (doi:10.1016/j.omto.2020.07.004)

46. Kaczmarczyk LS, Levi N, Segal T, Salmon-Divon M, Gerlitz G. 2022 CTCF supports preferentially short lamina-associated domains. Chromosome Res 30, 123–136. (doi:10.1007/s10577-022-09686-5)

47. Kamisuki S et al. 2009 A Small Molecule That Blocks Fat Synthesis By Inhibiting the Activation of SREBP. Chemistry & Biology 16, 882–892. (doi:10.1016/j.chembiol.2009.07.007)

48. DeBose-Boyd RA, Ye J. 2018 SREBPs in Lipid Metabolism, Insulin Signaling, and Beyond. Trends in Biochemical Sciences 43, 358–368. (doi:10.1016/j.tibs.2018.01.005)

49. Maja M, Tyteca D. 2022 Alteration of cholesterol distribution at the plasma membrane of cancer cells: From evidence to pathophysiological implication and promising therapy strategy. Front. Physiol. 13, 999883. (doi:10.3389/fphys.2022.999883)

50. Yan H, Tang G, Wang H, Hao L, He T, Sun X, Ting AH, Deng A, Sun S. 2016 DNA methylation reactivates GAD1 expression in cancer by preventing CTCF-mediated polycomb repressive complex 2 recruitment. Oncogene 35, 3995–4008. (doi:10.1038/onc.2015.423)

51. Gales L et al. 2022 Antidiabetics, Anthelmintics, Statins, and Beta-Blockers as Co-Adjuvant Drugs in Cancer Therapy. Medicina 58, 1239. (doi:10.3390/medicina58091239)

52. Snaebjornsson MT, Janaki-Raman S, Schulze A. 2020 Greasing the Wheels of the Cancer Machine: The Role of Lipid Metabolism in Cancer. Cell Metabolism 31, 62–76. (doi:10.1016/j.cmet.2019.11.010)

53. Nora EP et al. 2020 Molecular basis of CTCF binding polarity in genome folding. Nat Commun 11, 5612. (doi:10.1038/s41467-020-19283-x)

54. Hashimshony T et al. 2016 CEL-Seq2: sensitive highly-multiplexed single-cell RNA-Seq. Genome Biol 17, 77. (doi:10.1186/s13059-016-0938-8)

55. Kim D, Langmead B, Salzberg SL. 2015 HISAT: a fast spliced aligner with low memory requirements. Nat Methods 12, 357–360. (doi:10.1038/nmeth.3317)

56. Kim D, Paggi JM, Park C, Bennett C, Salzberg SL. 2019 Graph-based genome alignment and genotyping with HISAT2 and HISAT-genotype. Nat Biotechnol 37, 907–915. (doi:10.1038/s41587-019-0201-4)

57. Liao Y, Smyth GK, Shi W. 2014 featureCounts: an efficient general purpose program for assigning sequence reads to genomic features. Bioinformatics 30, 923–930. (doi:10.1093/bioinformatics/btt656)

58. Robinson MD, McCarthy DJ, Smyth GK. 2010 edgeR : a Bioconductor package for differential expression analysis of digital gene expression data. Bioinformatics 26, 139–140. (doi:10.1093/bioinformatics/btp616)

59. Ritchie ME, Phipson B, Wu D, Hu Y, Law CW, Shi W, Smyth GK. 2015 limma powers differential expression analyses for RNA-sequencing and microarray studies. Nucleic Acids Research 43, e47–e47. (doi:10.1093/nar/gkv007)

60. Reimers M, Carey VJ. 2006 [8] Bioconductor: An Open Source Framework for Bioinformatics and Computational Biology. In Methods in Enzymology, pp. 119–134. Elsevier. (doi:10.1016/S0076-6879(06)11008-3)

61. Law CW, Chen Y, Shi W, Smyth GK. 2014 voom: precision weights unlock linear model analysis tools for RNA-seq read counts. Genome Biol 15, R29. (doi:10.1186/gb-2014-15-2-r29)

